# Altered Copper Transport in Oxidative Stress-Dependent Brain Endothelial Barrier Dysfunction Associated with Alzheimer’s Disease

**DOI:** 10.1101/2024.08.28.610108

**Authors:** Md. Selim Hossain, Archita Das, Ashiq M. Rafiq, Ferenc Deák, Zsolt Bagi, Rashelle Outlaw, Varadarajan Sudhahar, Mai Yamamoto, Jack H. Kaplan, Masuko Ushio-Fukai, Tohru Fukai

## Abstract

Oxidative stress and blood-brain barrier (BBB) disruption due to brain endothelial barrier dysfunction contribute to Alzheimer’s Disease (AD), which is characterized by beta-amyloid (Aβ) accumulation in senile plaques. Copper (Cu) is implicated in AD pathology and its levels are tightly controlled by several Cu transport proteins. However, their expression and role in AD, particularly in relation to brain endothelial barrier function remains unclear. In this study, we examined the expression of Cu transport proteins in the brains of AD mouse models as well as their involvement in Aβ42-induced brain endothelial barrier dysfunction. We found that the Cu uptake transporter CTR1 was upregulated, while the Cu exporter ATP7A and/or ATP7B were downregulated in the hippocampus of AD mouse models, and in Aβ42-treated human brain microvascular endothelial cells (hBMECs). In the 5xFAD AD mouse model, Cu levels (assessed by ICP-MS) were elevated in the hippocampus. Moreover, Aβ42-induced reactive oxygen species (ROS) production, ROS-dependent loss in barrier function in hBMEC (measured by transendothelial electrical resistance), and tyrosine phosphorylation of VE-cadherin were all inhibited by either a membrane permeable Cu chelator or by knocking down CTR1 expression. These findings suggest that dysregulated expression of Cu transport proteins may lead to intracellular Cu accumulation in the AD brain, and that Aβ42 promotes ROS-dependent brain endothelial barrier dysfunction and VE-Cadherin phosphorylation in a CTR1-Cu-dependent manner. Our study uncovers the critical role of Cu transport proteins in oxidative stress-related loss of BBB integrity in AD.

**Highlights:**

1. Upregulation of the Cu importer CTR1 and downregulation of the Cu exporter ATP7A in the hippocampus of AD mouse models
2. Aβ42 increases CTR1 expression while reduces ATP7A and ATP7B levels in human brain microvascular ECs.
3. Aβ42 triggers increased reactive oxygen species (ROS) production in human brain microvascular ECs through a CTR1- and Cu-dependent manner.
4. Aβ42 induces endothelial barrier dysfunction in human brain microvascular ECs through a CTR1-Cu-ROS-pendent manner.

## 1. Introduction

Alzheimer’s disease (AD) is a multifactorial, progressive neurodegenerative disease characterized by the accumulation of beta-amyloid 40 (Aβ40) and 42 (Aβ42) [1, 2]. Although both Aβ42 and Aβ40 are generated from amyloid precursor protein (*APP*) through sequential proteolytic cleavages by β-secretase and γ-secretase [3], Aβ42 is more hydrophobic and prone to aggregation due to additional hydrophobic amino acids at its C-terminus, compared with Aβ40 [4]. The Aβ42/Aβ40 ratio in blood plasma serves as a strong biomarker for AD and cerebral amyloid angiopathy (perivascular deposition of Aβ) [5], another frequently observed pathological hallmark of AD [6–8]. Elevated expression of APP in cerebral endothelial cells (ECs) suggests pathological cerebrovascular reactivity to Aβ42 [9]. Additionally, inflammatory markers such as VCAM-1, ICAM-1, E-selectin, MCP-1, and TNFα are elevated in cerebrovascular tissues, exerting toxic effects on neurons in AD [10, 11]. Thus, vascular, especially EC dysfunction, is closely linked to AD. In AD, cerebrovascular abnormalities, particularly blood-brain-barrier (BBB) dysfunction, are associated with inflammation, altered cerebral blood flow, and hypoperfusion. Brain endothelial cells (BECs), forming the main component of the BBB, are connected by adherence and tight junctions along with pericyte and astroglia layers [12, 13]. Aβ42 induces oxidative stress, which plays a pivotal role in the pathogenesis in AD, including BBB dysfunction [14, 15]. NADPH oxidases (NOXes), key sources of reactive oxygen species (ROS), significantly contribute to BEC barrier dysfunction in AD [16, 17]. However, the mechanisms underlying ROS-dependent BBB disruption in AD remain elusive.

Copper (Cu) is an essential micronutrient that functions as an enzyme cofactor in various cellular process, including respiration, neurotransmitter synthesis, and antioxidant defense [18]. It also plays a role in intracellular signaling, synaptic transmission, catecholamine balance, and myelination in the central nervous system [19]. The brain has a high demand for Cu, with levels varying across different regions [20].The dentated gyrus molecular layer of the hippocampus, locus coeruleus, and subventricular zone in ventricles have higher Cu content than other brain regions [21, 22]. Cu is thought to enter the brain through BBB from the peripheral circulation and exit via the cerebrospinal fluid barrier [23]. In AD, Cu contributes to oxidative stress, tau protein hyperphosphorylation, and Aβ aggregation [24, 25]. AD patients often show higher Cu levels in serum and amyloid plaque [26, 27] [28–31]. Since excess Cu is toxic, Cu balance is crucial for proper brain function. This balance is maintained by Cu uptake transporter 1 (CTR1), Cu chaperons including Atox1, and Cu transporting/exporting ATPases (ATP7A and ATP7B) [32]. CTR1 has been found to be overexpressed in the brain barrier of North Ronaldsay sheep, facilitating Cu uptake, which is linked to neurogenerative disease [33]. Our research demonstrated that endothelial CTR1 promotes VEGF signaling and angiogenesis both in vitro and in vivo [34]. Furthermore, we showed that CTR1 expression is upregulated in vascular cells through HIF1α-dependent pathway under hypoxic conditions [34]. ATP7A plays a vital role in providing Cu to key enzymes involved in neurotransmitter synthesis [35] and neuronal function [22] as well as in reducing intracellular Cu levels. A reduction in ATP7A can lead to significant neurological effects. Mutations in ATP7A or ATP7B cause Menkes and Wilson diseases, both of which involve neurological disorders [32]. Additionally, variations in the ATP7B gene are observed in AD patients [36], and AD mice models have demonstrated disrupted microglial Cu trafficking [37]. However, the role of Cu and Cu transport proteins in Aβ42-induced, ROS-dependent dysfunction of the brain endothelial cell barrier remains unknown.

In this study, we investigated the expression of Cu transport proteins in AD and identified similar dysregulation (CTR1 upregulation and ATP7A downregulation) in mice AD models, suggesting a disruption of intracellular Cu homeostasis in AD. We further explored the role of Cu and CTR1 in the loss of endothelial barrier integrity in Aβ42-stimulated hBMECs. Our findings indicate that Aβ42 induces ROS production, resulting in the phosphorylation of VE-cadherin and increased endothelial permeability. This is the first report demonstrating the role of Cu transport proteins in Aβ42-induced brain endothelial barrier dysfunction, highlighting the significance of Cu transport dysregulation in oxidative stress-related loss of endothelial barrier integrity in AD. Therefore, regulating Cu homeostasis may represent an important therapeutic strategy for treatment of AD pathology.

## 2. Materials and Methods

### 2.1 Animals

24 months old male/female triple transgenic AD (3x Tg-AD) mice harboring *APPswe, PSEN1M146V* and *tauP301L* transgenes, 6 months old male 5x-FAD mice having 5 AD linked mutations (APP K670_M671delinsNL (Swedish), APP I716V (Florida), APP V717I (London), PSEN1 M146L (A>C), PSEN1 L286V) [38, 39] and age-matched non-transgenic mice with the same genetic background as controls were used for our study. All mice were maintained at the animal facilities of Augusta University. All studies were carried out in accordance with the guidelines of approval Committee at Medical College of Georgia, Augusta University.

### 2.2. Aβ42 peptide purification and preparation

The Aβ42 peptide having sequence DAEFRHDSGYEVHHQKLVFFAEDVGSNKGAIIGL MVGGVVIA used in this study was purified from pET28b-GroeES-ubiquitin-Aβ42 construct by using previously described methods [40]. The sense and antisense primers for polymerase chain reaction (PCR) of ubiquitin-Aβ42 is: (sense 5’-CGC-GGA-TCC-CAG-ATC-TTT-GTG-AAG-AC-3’ ; antisense 5’-CCG-CTC-GAG-TCA-CGC-TAT-GAC-AAC-ACC-GCC-3’. Briefly, inclusion bodies were recovered from fusion protein produced in E. Coli. To remove GroeES-ubiquitin, fusion proteins were digested with Usp2-cc as deubiquitylating enzyme [41] followed by methanol precipitation and the desired peptide was collected after centrifugation at 2000 × g for 10 min. Purified peptide was dissolved in 0.1 % NH_4_OH with the concentration of 2 mg/ml. During experiments, peptide was diluted with cell culture media to the desired concentration.

### 2.3. Scrambled Aβ42 preparation

Scrambled Aβ42 (rPeptide, GA, USA) having sequence KVKGLIDGAHIGDLVYEFMDS NSAIFREGVGAGHVHVAQVEF was used as negative control which does not aggregate as native Aβ42. Peptide was dissolved in 0.1 % NH_4_OH with the concentration of 2 mg/ml. During experiments, peptide was diluted to the desired concentration with cell culture media.

### 2.4. Cell culture

Human Brain Microvascular Endothelial Cells (hBMEC) (Science Cell Research Laboratories) were grown on 0.1 % gelatin coating in Endothelial cell medium supplemented with Cell Growth supplement, Fetal Bovine Serum (FBS) and penicillin/streptomycin solution having passage number 3-5.

### 2.5. Transfections

Control siRNA and CTR1 siRNA were obtained from Ambion and Invitrogen respectively. HBMEC was seeded into 100 mm culture dishes 1 day before transfection. Oligofectamine (Invitrogen) was used for the transfection following the manufacturer’s procedure. 30 nM siRNAs were used for the transfection. After transfection, media was changed to growth medium and incubated for 48 h at 37 ^⁰^C before performing experiments.

### 2.6. Immunoblotting

Human brain tissues, hippocampus and cortex tissues isolated from mouse brain were homogenized in RIPA buffer (5 mM Tris-HCl (pH 7.6), 150 mM NaCl, 1 % Triton X-100, 1 % Sodium deoxycholate, 0.1 % SDS) with protease inhibitor followed by 30 min incubation in rocker as described before [42]. For in vitro experiments, cells were lysed using the same RIPA buffer. After measurement of protein concentration using Bio-Rad protein assay dye, equal amounts of proteins were separated by SDS-PAGE and transferred onto nitrocellulose membrane. The membrane was then incubated with primary antibodies overnight followed by secondary antibodies incubation for 1 h to visualize protein expression by ECL (Amersham). Primary antibodies used for this study are anti-CTR1 (Abcam, 1: 2000) for human samples, anti-CTR1 (home-made, 1: 1000) for mouse samples, anti-Atox1 (home-made, 1: 1000), anti-ATP7A (LS Bioscience, 1: 1000), anti-ATP7B (Santa Cruz, 1: 1000), anti-ATP7B (Gene Tax, 1: 1000) for mouse samples, anti-APP (Invitrogen, 1: 1000), anti-phospho-VE-cadherin (Y658) (Invitrogen, 1 : 1000), total anti-VE-cadherin (Santa Cruz, 1 : 1000). Goat anti-rabbit IgG-HRP conjugate (Bio-Rad, 1: 2000) and Goat anti-mouse IgG-HRP (Bio-Rad, 1: 2000) were used as secondary antibodies. Image J was applied to quantify the band density.

### 2.7. Immunohistochemistry

Frozen sections were prepared by OCT embedding, 7 μm thick sections following overnight 4% PFA incubation and sucrose dehydration, then stained with hematoxyline-eosin (H/E). Counterstaining with hematoxylin was performed. For the CTR1 immunostaining, CTR1 (Abcam) primary antibody and Alexa Fluor 488 conjugated goat anti-rabbit IgG (Invitrogen) secondary antibody was used. Nuclei were stained with DAPI (Vector Laboratories), and images were captured by fluorescence microscopy (Keyence, BZ-X700, Osaka, Japan).

### 2.8. Quantitative real time polymerase chain reaction (qRT-PCR)

Total RNA was isolated from mouse hippocampus and cortex tissues and HBMEC using TRI reagent (Molecular Research Center, Inc.). 2 μg of total RNA was reverse transcribed to make cDNA by using a cDNA reverse transcription kit (Applied Biosystem). The PCR was performed following manufacturer’s procedures using ABI prism 7000 detection system (Applied Biosystems) and QuantiFast SYBR Green PCR kit (Qiagen). Gene expressions were normalized by 18s rRNA (internal control for human mRNA) and HPRT (internal control for mouse mRNA) respectively. Primer sequence for qPCR is: Human CTR1: Forward-5’-CCA-GGACCAAATGGAA-CCA-TCC-3’ and Reverse-5’-ATA-CCA-CAG-CCT-GGC-ACA-ACCT-3’; Mouse ATP7A: Primers purchased from Qiagen with Cat # Mm_Atp7a_1_SG QuantiTect Primer Assay (QT00152677) (sequence not available); Human ATP7B: Forward-5’-GGA-CCA-CAACATCATTCCAGGAC-3’ and Reverse-5’-ATG-AGC-ACG-TCCATGTTGGCTG-3’;Mouse ATP7B: Forward-5’-ATC-ATC-CCA-GGA-CTG-TCCGTTC-3’ and Reverse-5’-ATG-TTG-GCG-GAC-CTGTGTCTCA-3’. Data were expressed as fold changes.

### 2.9. Transendothelial Electrical Resistance (TEER) to measure a barrier function in hBME

The TEER of ECs with or without Aβ42 treatment was measured using electric cell-substrate sensing (ECIS) in an ECIS system Model 1600R (Applied Biophysics) [43]. Briefly, transwells were first coated with 0.1 % gelatin and equal number of cells (35 × 10^3^/well) were seeded on 8W10E+ PET arrays and resistance values of the cells were > 1000 Ω of steady state at a frequency of 4000 Hz. Data obtained as resistance values were plotted as normalized resistance.

### 2.10. Measurement of ROS

After treatment, cells were incubated with 20 μM H2DCFDA (5,6-chloromethyl-2’,7’-dichlorodihydro fluorescein diacetate, acetyl ester, Invitrogen) for 6 min at 37 ⁰C as described [44]. The cells were then mounted using VECTASHIELD mounting medium with 4,6-diamidino-2-phenylindole (DAPI) (Vectorlabs). The DCF fluorescence intensity in cells was measured by fluorescence microscopy (Keyence, BZ-X700, Osaka, Japan) and quantified using Image J.

### 2.11. Measurement of metal contents by Inductively coupled mass spectrometry (ICP-MS)

Brain tissues were digested by adding 100 μl concentrated HNO3 (trace metal grade, Fisher) and heating at 90°C for 1 hour in a heating block. Retinas were digested by adding 50 μl concentrated HNO3 (trace metal grade, Fisher) and heating at 90°C for 1 hour in a heating block. Brain tissue samples were diluted to a total 1.5ml with 1% HNO3, and retina were diluted to a total 1.4 ml with 1% HNO3. Samples were then diluted with 1% nitric acid. A NIST SRM 1683f was prepared to ensure accuracy of the calibration curve. ICP-MS was performed using an Agilent 8900 triple quad equipped with an SPS autosampler. The system was operated at a radio frequency power of 1550 W. Metal contents in the brain tissues and retinas were calculated from calibration curves as previously described [45].

### 2.12. Statistical Analysis

Results are expressed as mean ± SEM. Statistical significance was evaluated by a two-tailed unpaired t-test or two-way Analysis of Variance (ANOVA) test, followed by Tukey post-hoc to specify the differences between the groups. Prism V10 (GraphPad Software) was used for the statistical test. A p value of *p<0.05, **p<0.01, #p<0.001, ##p<0.0001 were considered statistically significant.

## 3. Results

### 3.1 Upregulation of CTR1 and Downregulation of ATP7A Expression in the Hippocampus of Mouse AD Models

The hippocampus is a crucial region for cognition and memory, and BBB disruption in the hippocampus is recognized as an early biomarker of cognitive dysfunction in AD [46]. Additionally, Cu homeostasis is vital for hippocampus function [47]. Therefore, we evaluated the expression of Cu transport proteins in the hippocampus of established 3xTg and 5xFAD mouse models of AD. As shown in Figures 1A and 1B, the expression of Cu uptake transporter CTR1 protein was significantly increased, whereas Cu exporter ATP7A protein expression was reduced in both AD models compared to control mice. Notably, the protein levels of Atox1 and ATP7B showed no significant differences between AD models and control groups. Furthermore, we verified the overexpression of amyloid precursor protein (APP) in the hippocampus of AD mice compared to control mice. Further analysis of mRNAs revealed a significant increase in CTR1 mRNA levels and a decrease in ATP7A mRNA levels, while the mRNA levels of ATP7B and Atox1 remained unchanged in both AD models compared to control mice (Figures 2A and 2B). We then analyzed the levels of Cu in the hippocampus of 5xFAD AD mouse models, along with iron (Fe) and zinc (Zn), which have also been implicated in AD pathology [31]. Since the retina is an extension of the central nervous system and is known to exhibit changes associated with brain pathology in AD [48, 49], we also examined levels of Cu, and other metals in the retina. Notably, ICP-MS analysis revealed increased levels of Cu, Fe, and Zn in the hippocampus, but not in the retina, of 5xFAD AD mice (Supplemental Figure 1). These findings indicate that the transcriptional upregulation of CTR1 and downregulation of ATP7A may contribute to elevated Cu levels in the hippocampus of AD mouse models.

**Figure 1.**
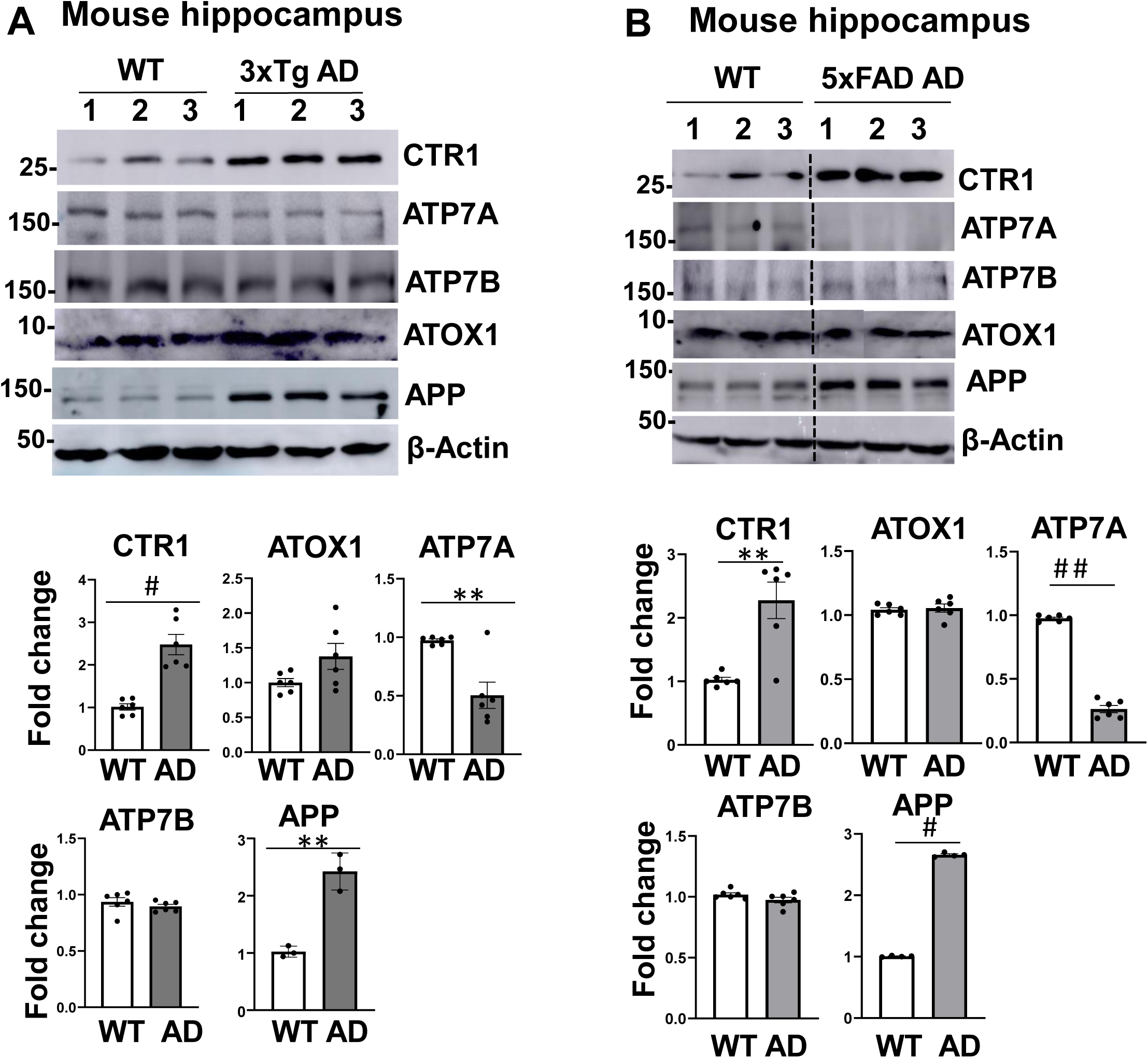
CTR1 protein expression is increased but ATP7A protein expression is decreased in hippocampus from mouse AD models. Protein expression for Cu transport proteins in hippocampus tissues isolated from WT and 3xTg AD mouse model (A) as well as WT and 5x FAD AD mouse model (B). Amyloid precursor protein (APP) expression was used as a positive control for AD phenotype. Data are mean ± SEM (n=5-6). **p<0.01, #p<0.001, ##p<0.0001.

**Figure 2.**
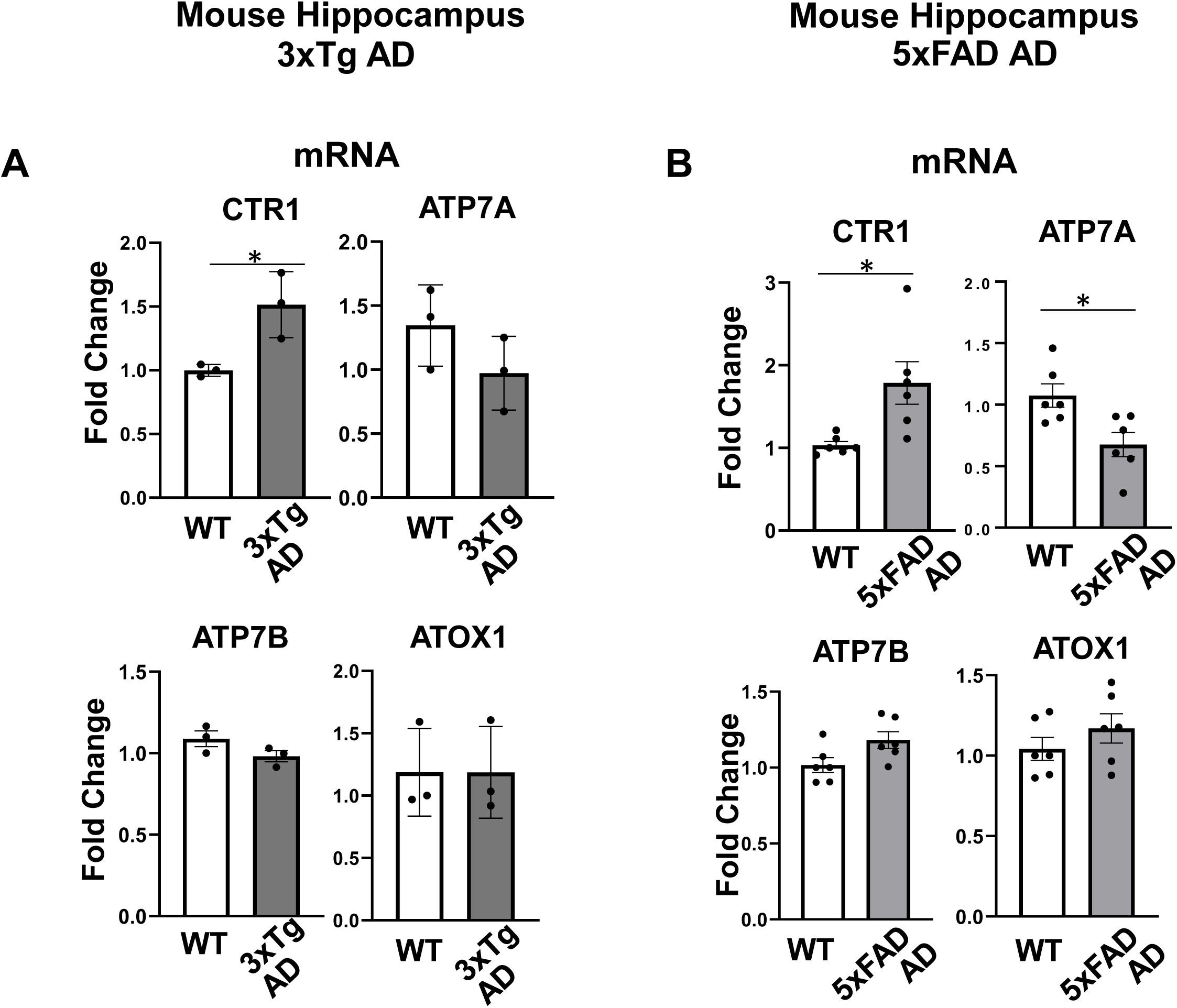
CTR1 mRNA expression is increased but ATP7A mRNA expression is decreased in hippocampus from mouse AD models. mRNA expression for Cu transport proteins in hippocampus tissues isolated from WT and 3xTg AD mouse model (A) as well as WT and 5x FAD AD mouse model (B). Data are mean ± SEM (n=3-6). *p<0.05.

### 3.3 Upregulation of CTR1 and Downregulation of ATP7A and ATP7B in Human Brain Microvascular Endothelial Cells (hBMECs) stimulated with Aβ42

We then investigated the effects of Aβ42 on the expression of Cu transport proteins in hBMECs. As shown in Figure 3, Aβ42 stimulation led to an increase in CTR1 protein expression in a dose- and time-dependent manner (Figures 3A and 3C). Concentrations higher than 20 µM and stimulation durations longer than 16 hours resulted in toxicity, as evidenced by trypan blue staining (data not shown). Conversely, Aβ42 stimulation decreased ATP7A and ATP7B protein expression in a dose- and time-dependent manner. Additionally, we confirmed that a scrambled Aβ42 control peptide, at the same concentration and incubation time as Aβ42, did not affect the expression of these Cu transport proteins in hBMECs (Figure 3B). Notably, mRNA expression analysis indicated that Aβ42 (20 μM) stimulation increased CTR1 mRNA levels and reduced ATP7A mRNA levels in a time-dependent manner, while the mRNA expression of Atox1 and ATP7B remained unchanged. These findings suggest that Aβ42 stimulation upregulates CTR1 expression at the transcriptional level and downregulates ATP7A expression at the transcriptional level and ATP7B expression at the post-translational level in hBMECs.

**Figure 3.**
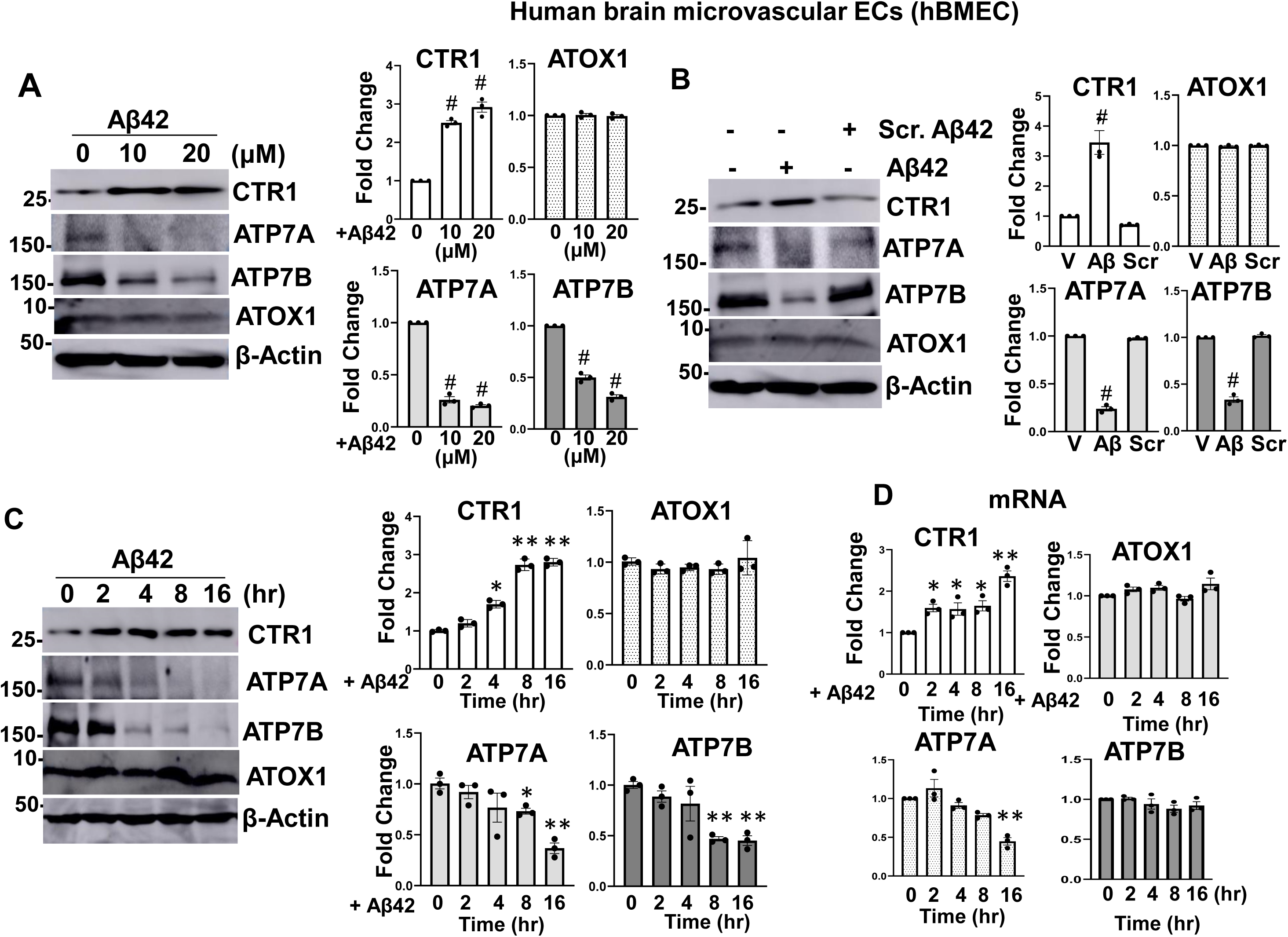
Aβ42 stimulation induces upregulation of CTR1 expression and downregulation of ATP7A and ATP7B expression in hBMEC. **A, B, C.** hBMECs were stimulated with Aβ42 at 0-20 μM for 16 h (A), or 20 μM Aβ42 or scrambled Aβ42 (Scr. Aβ42) for 16 hr (B), or 20 μM Aβ42 for indicated time (0-16 hr)(C). These lysates were used for western blotting to measure protein expression of Cu transport proteins. **D.** hBMEC stimulated with 20 μM Aβ42 for indicated time were used to measure Cu transport protein mRNA using qPCR. Data are mean ± SEM (n=3). *p<0.05. **p<0.01, #p<0.001.

### 3.4 Aβ42 induces endothelial barrier dysfunction in a CTR1-Cu-ROS-dependent manner in hBMECs

Since CTR1 was markedly upregulated in the hippocampus of mouse AD models as well as in hBMECs stimulated with Aβ42, we investigated the role of Cu and CTR1 in Aβ42-induced brain EC barrier dysfunction, which is reported to be ROS-dependent [17]. A transendothelial electrical resistance (TEER) assay revealed that in a confluent monolayer of ECs, Aβ42 stimulation reduced electrical impedance due to the loss of intercellular junction and barrier function, in a time-dependent manner (Figures 4A and 4B). Notably, pretreatment of hBMECs with either the cell-permeable Cu chelator, Tetrathiomolybdate (TTM) or the thiol antioxidant N-acetylcysteine (NAC)(Figure 4A), or transfected with CTR1 siRNA (Figure 4B), completely prevented Aβ42-induced EC barrier dysfunction. These results suggest that Aβ42-induced disruption of endothelial barrier integrity is dependent on CTR1, Cu, and ROS.

**Figure 4.**
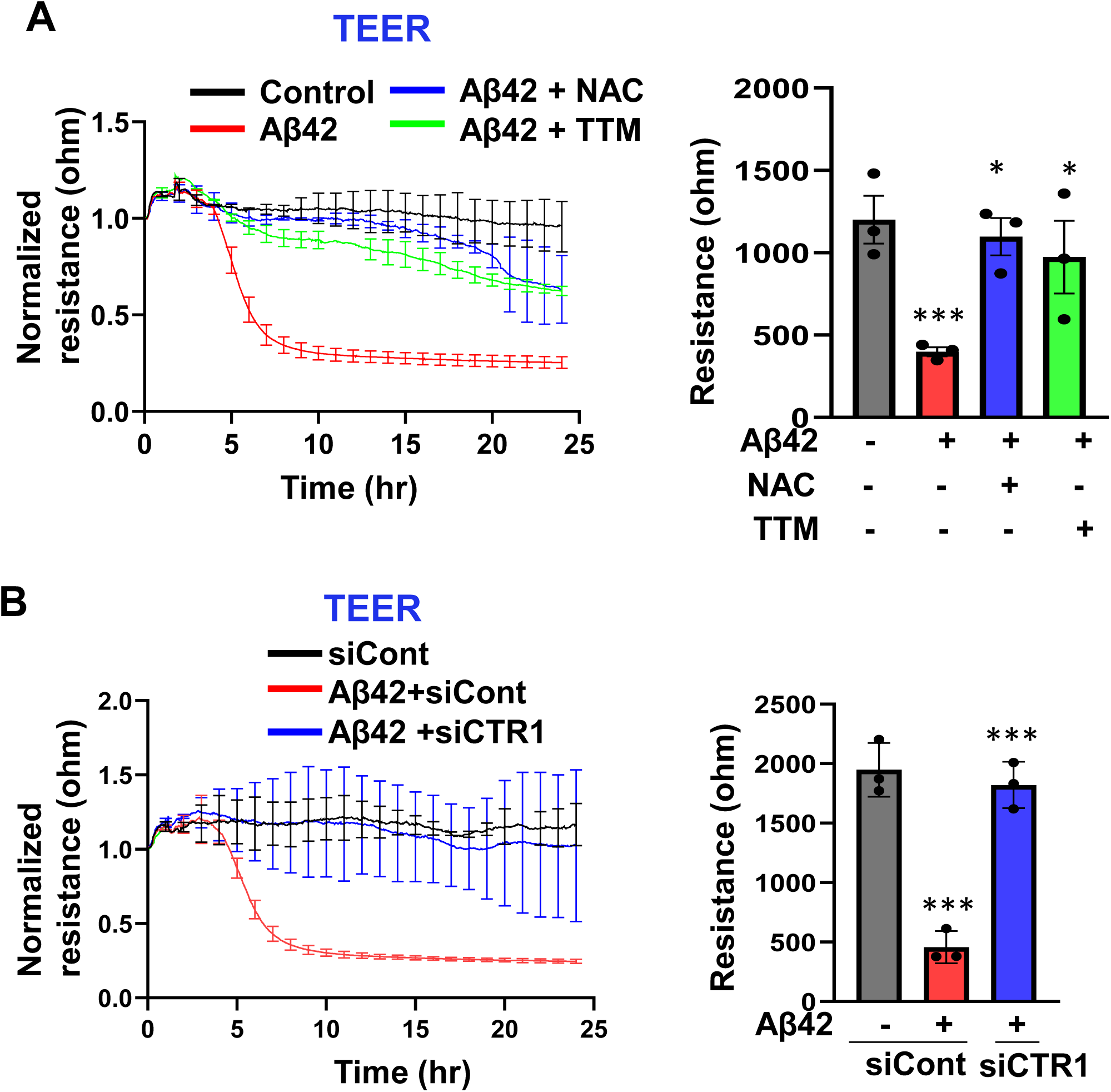
Aβ42-induces endothelial barrier dysfunction in a CTR1-Cu-ROS-pendent manner in hBMECs. **A and B**. hBMEC were pretreated with thiol antioxidant N-Acetyl Cysteine (NAC)(10 mM) for 1 h or Cu chelator, TTM (20 nM) for 24 hr (A), or transfected with siRNAs for CTR1 or scrambled control (30 nM)(B). These ECs stimulated with 20 μM Aβ42 for indicated times were used for TEER assay to measure EC permeability. Data analysis are shown in the right panel.

We next examined the relationships between the CTR1-Cu axis and ROS production induced by Aβ42 in hBMECs. We found that Aβ42, but not scrambled Aβ42, increased intracellular redox status detected by 2’,7’-dichlorodihydrofluorescein diacetate (DCF-DA) in a time-dependent manner, which was abolished by the overexpression of catalase (Figures 5A and 5B). This Aβ42-induced H_2_O_2_ production was inhibited by the Cu chelator TTM or CTR1 siRNA (Figures 5C and 5D). These results suggest that Aβ42 induces ROS production in a CTR1-Cu-dependent manner, which promotes endothelial barrier dysfunction in hBMECs.

**Figure 5.**
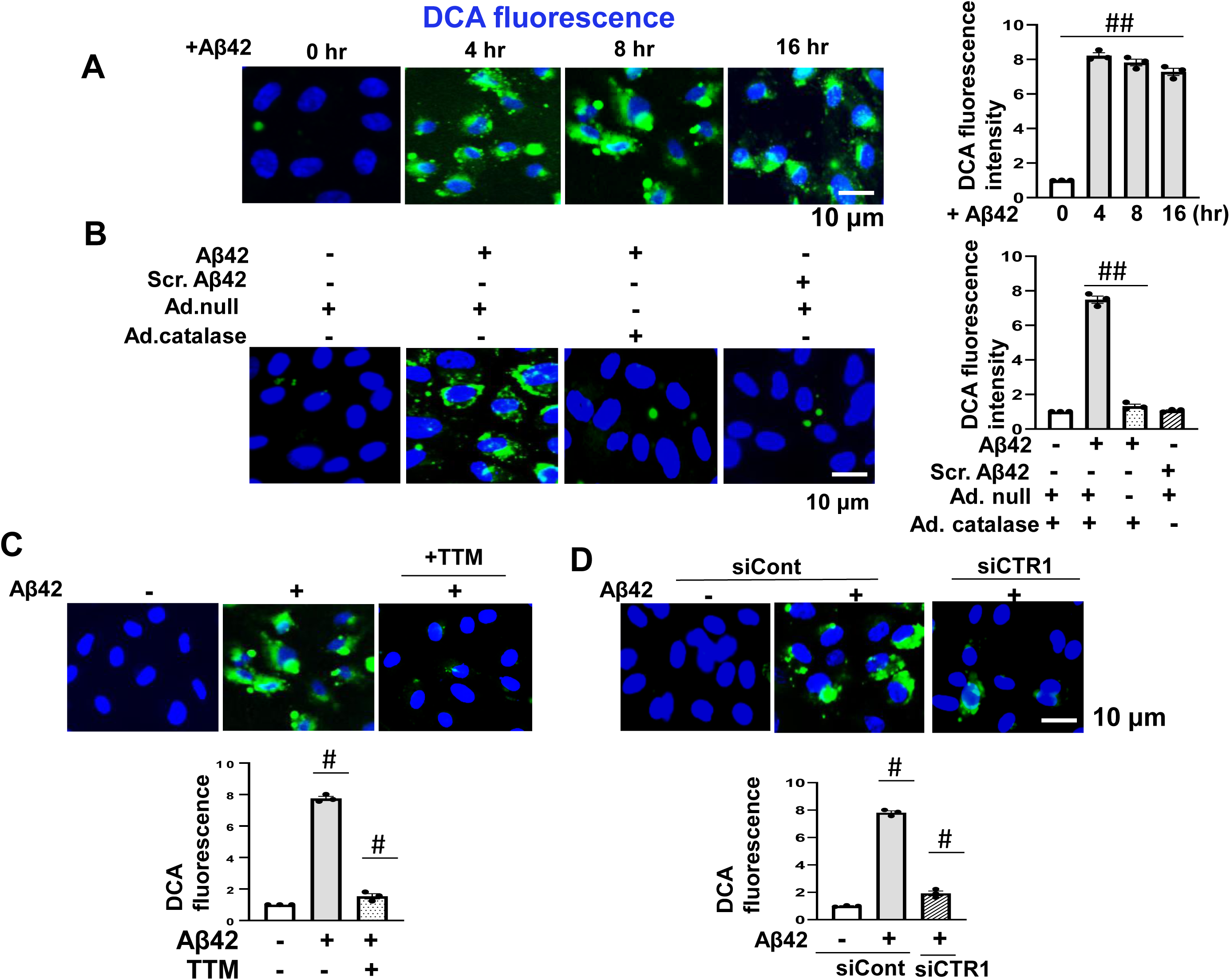
Aβ42 stimulation increases ROS production in a CTR1-Cu dependent manner in hBMECs. **A and B,** hBMECs stimulated with Aβ42 (20 μM) for the indicated time (0-16hr) (A), or transduced with adenovirus expressing catalase (Ad. Catalase) stimulated with Aβ42 (20 μM) for 16hr, or exposed to scrambled Aβ 42 (Scr. Aβ42) (20 μM) for 4 hr were used to measure DCF fluorescence (ROS production) (**B**). Representative images for DCF fluorescence with DAPI (blue) staining (Left) and quantification of DCF fluorescence intensity expressed as a fold change from the basal control (Right) are shown. Data are mean ± SEM (n=3). ****p<0.0001. **C and D,** hBMECs pretreated with TTM (20 nM) for 24 h (**C**), or transfected with siRNAs for CTR1 or control (30 nM) (**D**) were stimulated with 20 μM Aβ42 for 4 hr. These ECs were used for DCF fluorescence assay to measure ROS production. Representative images for DCF fluorescence with DAPI (blue) staining (Left) and quantification of DCF fluorescence intensity expressed as a fold change from the basal control (Right) are shown. Data are mean ± SEM (n=3) #p<0.001.

### 3.5 Aβ42 induces phosphorylation of VE-Cadherin in a CTR1-Cu-ROS-dependent manner in hBMECs

We next investigated the molecular mechanisms underlying Aβ42-induced loss of endothelial barrier function in hBMECs. It has been reported that Aβ42-induced ROS are required for the tyrosine phosphorylation of the adherence junction protein VE-Cadherin, which promotes brain ECs permeability [17]. We thus examined the roles of CTR1 and Cu in this response. We found that treatment with Aβ42 in hBMECs induced tyrosine phosphorylation of VE-Cadherin, which was almost completely inhibited by the Cu chelator TTM, the antioxidant NAC (Figure 6A), or CTR1 siRNA (Figure 6B). Notably, scrambled Aβ42 peptide had no effect on this response (Figure 6C). These results suggest that Aβ42 induces phosphorylation of VE-Cadherin in a CTR1-Cu-ROS-dependent manner in hBMECs, which may contribute to EC barrier dysfunction.

**Figure 6.**
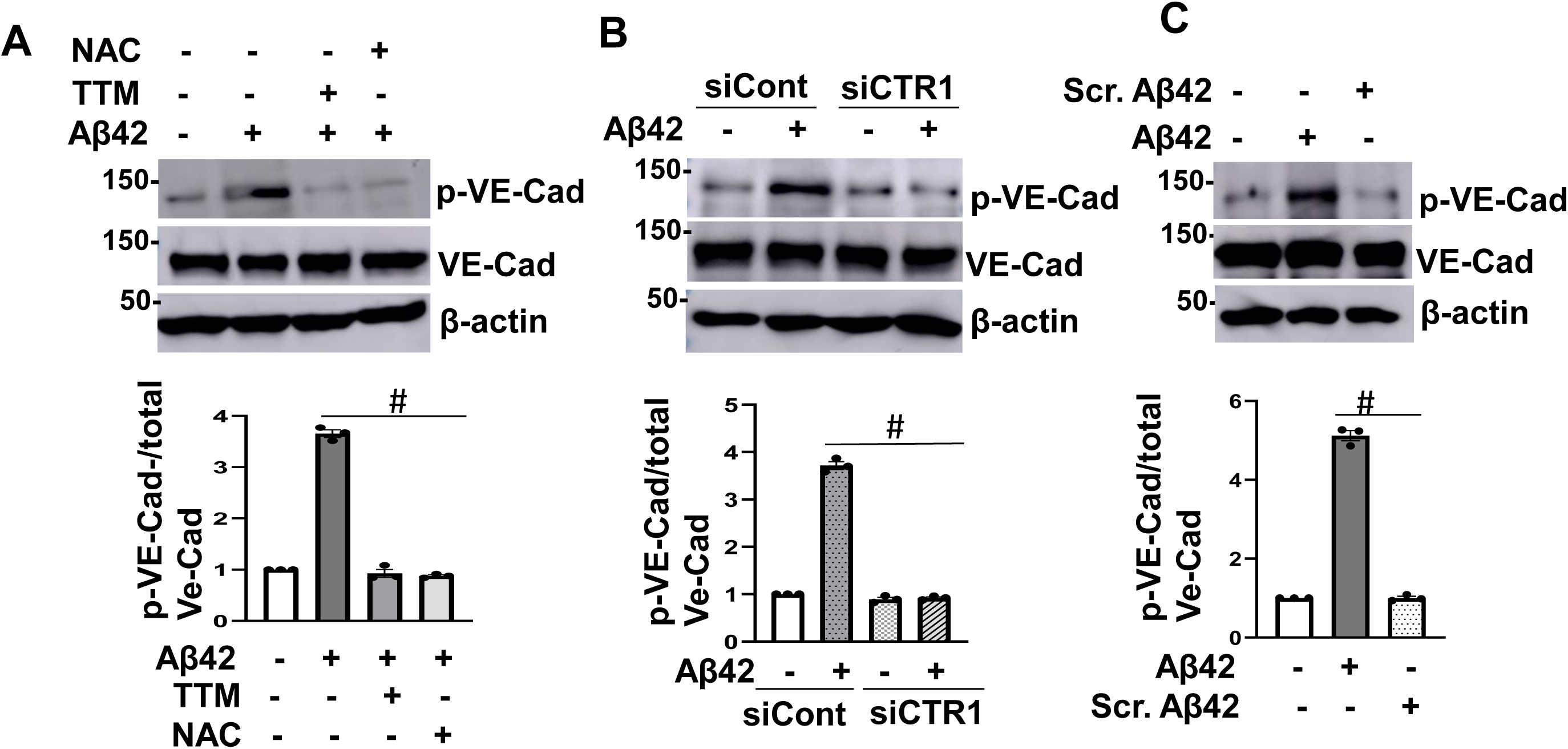
Aβ42-induced tyrosine phosphorylation of VE-cadherin in hBMECs is dependent on CTR1, Cu, and ROS. **A,B,C.** hBMECs pretreated with TTM (20 nM for 24 h) or antioxidant NAC (10 mM for 1 h)(**A**), or transfected with siRNAs for CTR1 or control (30 nM)(**B**) were stimulated with Aβ42. In parallel, hBMECs were stimulated with scrambled Aβ42 (20 μM) (**C**). Lysates from these cells were used for western blotting with anti-VE-Cadherin (VE-Cad)-pY658 or anti-VE-Cad or β-actin (loading control) antibodies. Bar graphs represent the fold changes from the basal control. Data are mean ± SEM (n=3). ***p<0.001, ****p<0.0001.

## 4. Discussion

Cu plays a crucial role in the pathology of AD [1–9]. The ECs of the BBB serve as a major gateway for Cu to enter the brain from the peripheral circulation, primarily through the Cu uptake transporter CTR1 [10], while the Cu exporter ATP7A facilitates Cu secretion from cells into the bloodstream [11]. However, the role of Cu transport proteins in AD pathogenesis, particularly their contribution to brain EC barrier dysfunction remains unclear. In this study, we investigated the changes in Cu transport protein expression in mouse AD brains, focusing on their implications for endothelial barrier dysfunction. Our findings reveal a significant upregulation of the Cu importer CTR1 and a concomitant downregulation of the Cu exporter ATP7A in the hippocampus of mouse AD models. In cultured hBMECs, we demonstrate that exposure to Aβ42 disrupts Cu homeostasis by upregulating CTR1 and downregulating ATP7A expression, leading to elevated ROS production. This process subsequently triggers the phosphorylation of VE-Cadherin, and stimulates brain endothelial barrier dysfunction, which may contribute to the loss of BBB integrity and progression of AD pathology (Figure 7). These findings suggest that dysregulation of Cu transport proteins play a critical role in AD pathophysiology, particularly brain endothelial barrier dysfunction.

**Figure 7.**
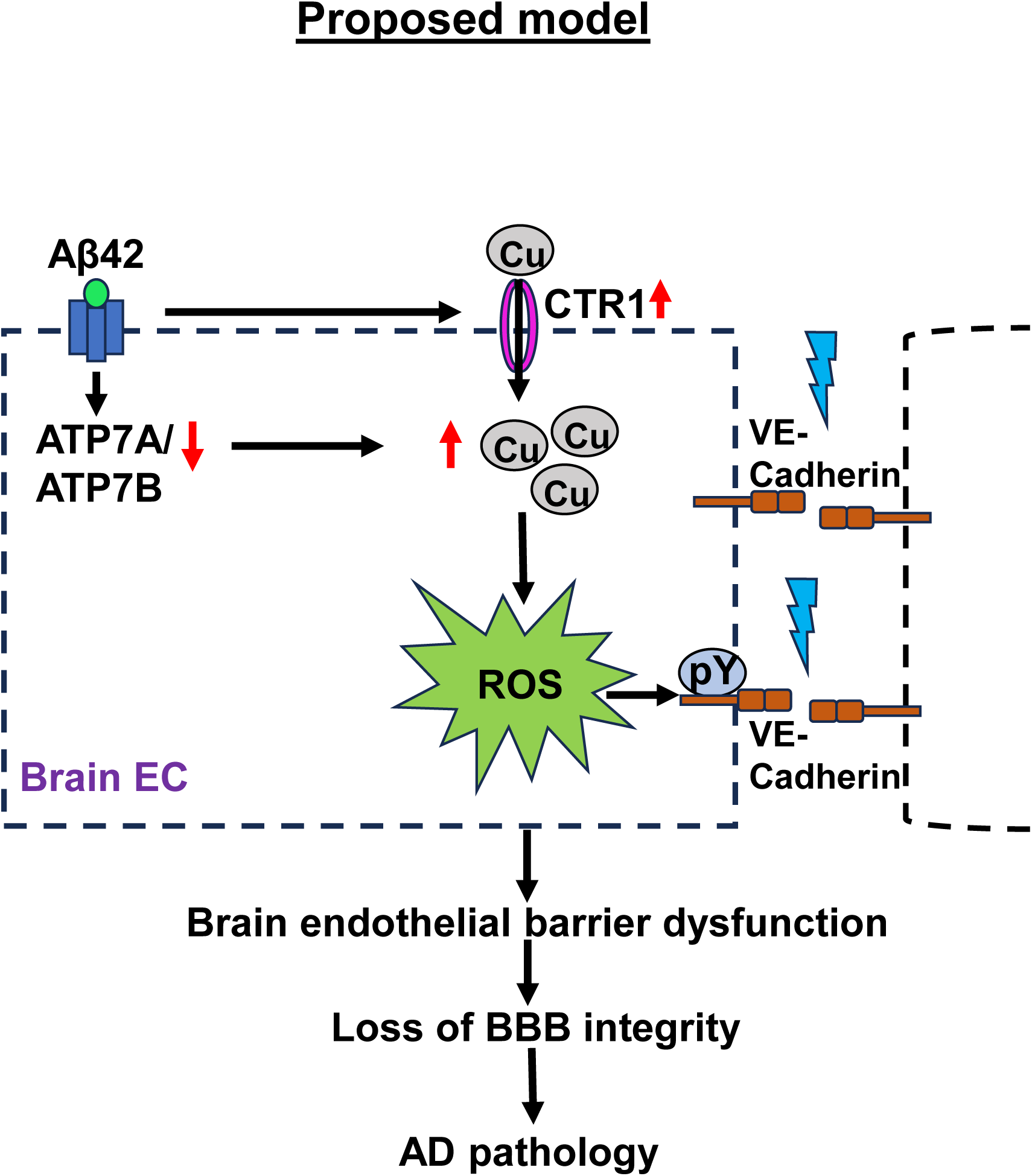
Schematic model: Aβ42 induces Cu dysregulation through upregulation of CTR1 and downregulation of ATP7A/ATP7B, which increases intracellular Cu and promotes ROS production and ROS-dependent tyrosine phosphorylation of VE-Cadherin, which causes brain endothelial barrier dysfunction, leading to loss of BBB integrity, resulting in progression of AD pathology.

The present study demonstrated increased expression of the Cu uptake transporter CTR1 and decreased expression of the Cu exporter ATP7A in the hippocampus of 3xTg and 5xFAD mouse models of AD. The hippocampus, a crucial region for cognition and memory, often exhibits early signs of AD. These findings suggest that elevated CTR1 expression leads to increased Cu influx, while reduced ATP7A expression would impair Cu export, resulting in Cu accumulation and subsequent toxicity. Notably, ATP7B protein remained unchanged in the hippocampus of mouse AD models compared to controls. This may indicate species-or brain region-specific regulation of ATP7B. Our results align with previous studies showing high concentrations of Cu in Aβ plaques, which are associated with plaque deposition in AD [95,96]. This dysregulation of Cu transport proteins could represent a potential mechanism through which Cu homeostasis is disrupted in AD, contributing to Cu accumulation in Aβ plaques and the progression of AD pathology.

The importance of Cu transporters, including CTR1 and ATP7A/B, in AD pathology is further supported by several pieces of evidence. Mutations in ATP7B (K832R), which impair ATP7B function, increase the risk for AD [12,13]. Notably, in an Aβ transgenic Drosophila model, reducing the Drosophila CTR1 homologue or overexpressing the Drosophila ATP7A/7B homologue reduces in vivo formation of Aβ oligomers, decreases oxidative stress, improves motor deficits, and prolongs lifespan [14], strongly implying an increase in intracellular Cu levels in the disease state as reported in our present studies. Cu transporting ATPases ATP7A and ATP7B play significant roles in the distribution of Cu in the central nervous system, and genetic mutations in these proteins lead to severe neurodegenerative states, as in Wilson and Menkes disease [13,15,16]. Additionally, we found that the expression levels of the Cu chaperone Atox1 did not differ significantly between AD models and control groups. Given that nuclear Atox1 functions as a Cu-dependent transcription factor under inflammatory conditions [17], further studies are needed to clarify its role in AD pathogenesis.

Our study also showed that CTR1 mRNA levels were significantly increased, while ATP7A mRNA levels were decreased, in the hippocampus of AD mouse model as well as in hBMEC stimulated with Aβ42, indicating transcriptional regulation of these proteins. The effects of Aβ42, a key pathogenic factor in AD, on Cu transport protein expression have not been reported in brain ECs. Our study revealed that Aβ42 stimulation, but not scrambled Aβ42, increased CTR1 protein and mRNA levels and decreased ATP7A protein and mRNA levels, as well as ATP7B protein without altering its mRNA, in a dose- and time-dependent manner. Several mechanisms for the regulation of CTR1 and ATP7A expression have been reported. For example, CTR1 expression is increased by transcription factors including metal-responsive transcription factor-1(MTF-1[19], specificity protein 1 (Sp1), or hypoxia inducible factor (HIF) 1 α or 2α [20–22]. ATP7A transcription is shown to be regulated by HIF2α [23].

BBB dysfunction has been implicated in the progression of neurodegeneration and dementia in AD by reducing the clearance of toxic metabolites, including Aβ, and by limiting the delivery of oxygen and nutrient to the brain. Elevated Cu levels have been found in amyloid plaques [24–26], which promotes Aβ peptide aggregation, neurotoxicity, and ROS production, leading to oxidative damage and BBB disruption, as evidenced by the increased permeability of brain ECs [8,9]. However, the role of Cu and Cu transport proteins in Aβ42-induced, ROS-dependent disruption of brain EC barrier has not been previously reported. Our findings demonstrate that the increase in Aβ42-induced EC barrier dysfunction is inhibited by the Cu chelator TTM, the antioxidant NAC, or knockdown of the Cu importer CTR1. Furthermore, Aβ42-induced ROS production was markedly reduced by Cu chelation, CTR1 siRNA, or catalase overexpression. These findings suggest that Aβ42 disrupts the brain EC barrier via a CTR1-Cu-ROS (H_2_O_2_)-dependent pathway. Previous studies have shown that Aβ42-induced ROS production is required for the tyrosine phosphorylation of VE-cadherin, which is crucial for the loss of junctional integrity and permeability of EC [27]. Our study, using the Cu chelator, antioxidant, and CTR1 siRNA, revealed that Aβ42, but not scrambled Aβ42, induces VE-Cadherin phosphorylation through a CTR1-Cu-ROS-dependent mechanism, thereby contributing to endothelial barrier dysfunction in AD. It is important to note that the BBB is maintained not only by adherens junctions but also by tight junctions (TJs). Indeed, Aβ42 stimulation has been shown to downregulate TJs proteins, such as occludin and ZO-1 and −2, in human brain ECs [16]. Therefore, future studies should explore the role of the CTR1-Cu axis in this response.

There are several possible mechanisms by which Aβ42 induces ROS production in a Cu-dependent manner. First, the Cu chaperone Atox1 may be involved, as we previously reported that Atox1 functions as a Cu dependent transcription factor, increasing the transcription of NOX organizer p47phox, leading to ROS production in ECs stimulated with inflammatory cytokine TNFα [28,29]. Although Nox1 and Nox2 have been proposed in Aβ induced BBB dysfunction [27,30], the role of Atox1 in those responses remains unclear. Second, Cu may enhance NADPH oxidase activity by upregulating phospholipase D [31]. Third, Cu can bind to Aβ42, contributing to ROS production [9]. Investigating the detailed mechanisms by which the CTR1-Cu axis promotes ROS production and its role in EC permeability, including its crosstalk with other metals, will be the focus of future research [18,32].

Our study demonstrated elevated Cu levels in the hippocampus, but not in the retina, of 5XFAD AD mouse models, as a result of the upregulation of CTR1 and the downregulation of ATP7A. Consistent with our results, previous reports show that amyloid plaques in both AD patients and AD mouse models are known to accumulate Cu [28–31], and that serum total or labile Cu levels are elevated in AD patients [26, 50, 51]. In contrast, Cu concentrations in the brain of AD patients and mouse models have been shown to be lower compared to healthy controls [52–54]. Mashkani et al. reported reduced Cu levels in both the hippocampus and retina of PP/PS1 AD mouse models [54]. The reason for this discrepancy remains unclear, but it may suggest that Cu homeostasis in AD is influenced by factors such as the location of Cu (intracellular vs. extracellular), the presence of labile Cu, specific brain regions, and disease progression [18]. Notably, we also observed increased levels of Zn and Fe in the hippocampus, but not in the retina, of 5XFAD AD mouse models, which aligns with other reports of metal dysregulation in AD brains [31]. These findings underline the importance of understanding the roles of Cu transporters and chaperones in AD pathology and their crosstalk with other metals, which could have significant implications for the developing new therapeutic strategies.

In summary, our findings advance the understanding of the role of Cu transport proteins in AD pathogenesis. Further research is needed to investigate the specific role of endothelial CTR1 in BBB dysfunction in vivo as well as the mechanisms by which Aβ42 induces CTR1 upregulation and ATP7A downregulation. Additionally, the effects of reduced ATP7A-mediated Cu accumulation on ROS-dependent endothelial barrier dysfunction warrant further exploration. Our studies suggest that targeting Cu transport proteins may offer new therapeutic strategies for mitigating various aspects of AD pathogenesis.

## Abbreviations

BBB: Blood-brain barrier
Aβ: beta-amyloid
AD: Alzheimer’s disease
hBMEC: Human brain microvascular endothelial cells
Cu: copper
CTR1: Cu uptake transporter 1
ROS: reactive oxygen species
NOXes: NADPH oxidases
TEER: transendothelial electrical resistance

## Acknowledgements

This work was supported by NIHR01HL160014 (to M.U.-F.), R01HL174014 (to T.F., M.U-F., V.S.), R01HL147550 (to T.F., M.U.-F.), P01HL160557 (to T.F, M.U-F), VA Merit Review grant I01BX001232 (to T.F.), AHA22TPA971863 (to T.F.), AHA21CDA941742 (to A.D.), NIH R01AG062655 (to FD) and SAGA23-1142437 award from Alzheimer’s Association (to FD), the National Institute on Aging (AG054651 to ZB. We thank Park Il Seon, Chosun University, Republic of Korea for providing Aβ42 and Usp2-cc plasmids. In addition, we acknowledge the assistance of Dr. Martina Ralle, Director of USR Elemental Analysis Core at OHSU (partial support from NIH (S10OD028492) for measuring metals levels using ICP-MS.

## Data availability

Date will be made available on request

## Supplementary Figure

**Supplementary Figure 1.**
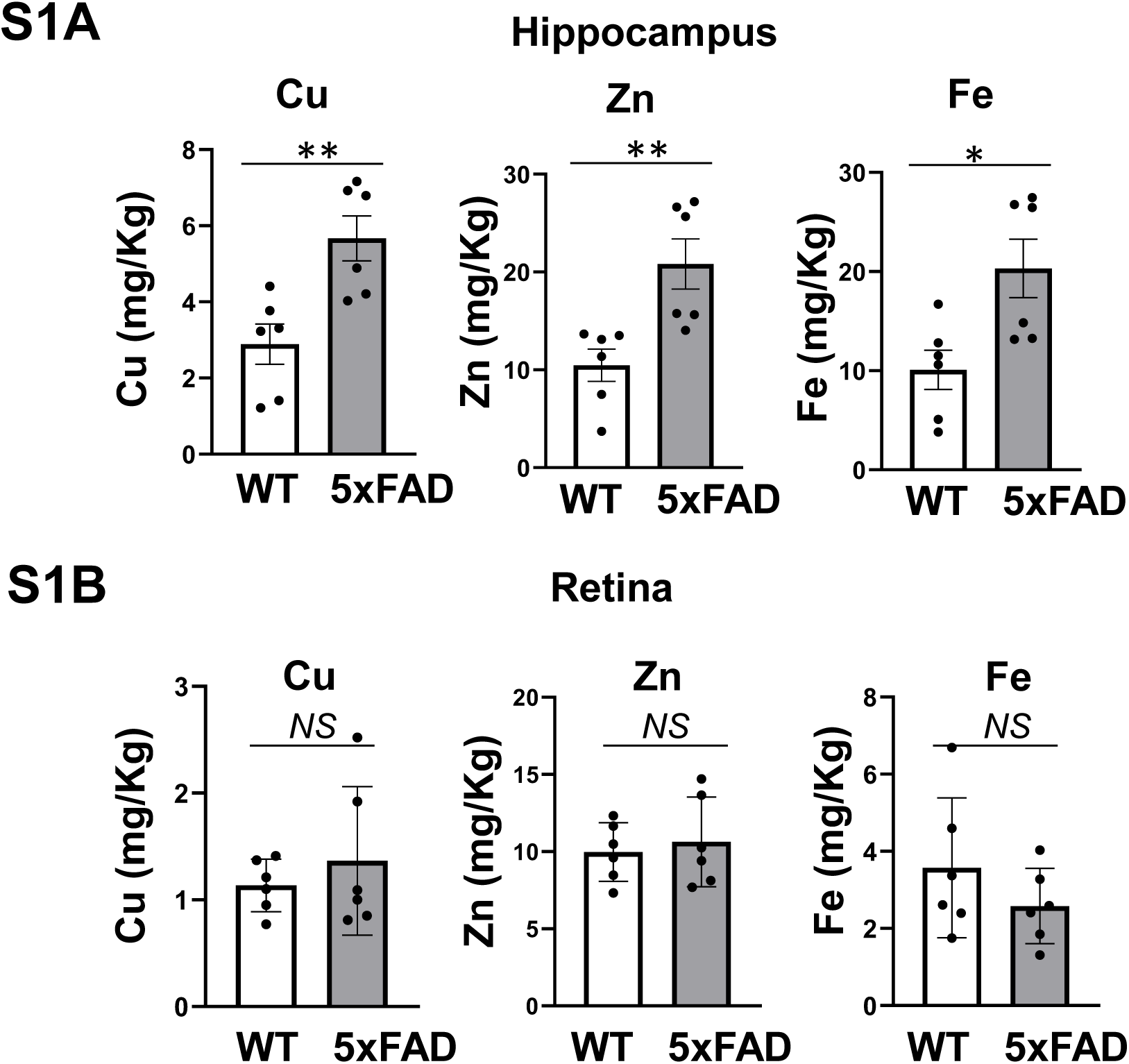
Cu levels are increased in the hippocampus, but not in the retina, from 5xFAD mouse AD model. Various metal ions including Cu levels were measured in hippocampus tissues (A) and retinal tissues (B) isolated from WT mice and 5xFAD AD mouse model using ICP-MS analysis. Data are mean ± SEM (n=6). *p<0.05, **p<0.01. NS: not significant.

